# Canonical Hedgehog Signaling Controls Astral Microtubules and Mitotic Spindle Orientation in Neural Progenitors and iPSCs

**DOI:** 10.1101/2025.02.23.639780

**Authors:** Fengming Liu, Anna Medyukhina, Kris M Olesen, Abbas Shirinifard, Hongjian Jin, Lei Li, Marina Mapelli, Khaled Khairy, Young-Goo Han

## Abstract

Mitotic spindle orientation is crucial for cell fate determination and tissue organization. Although the intracellular machinery governing spindle orientation is well characterized, whether and how secreted factors, such as morphogens, regulate this process remains poorly understood. This study investigated the role of Hedgehog (HH) signaling in modulating mitotic spindle orientation in neural progenitor cells and in induced pluripotent stem cells (iPSCs). Time-lapse microscopy of cerebral organoids and iPSCs revealed that HH signaling increases the angle of the mitotic spindle relative to the apical surface, prolongs mitosis, and enhances spindle rotation. Mechanistically, HH signaling reduces both the number and the length of astral microtubules, key regulators of spindle orientation. This reduction correlates with increased spindle angle in iPSCs. Furthermore, we show that canonical HH signaling, involving GLI-dependent transcriptional regulation, contributes to these effects. RNA sequencing and gene set enrichment analysis (GSEA) revealed that HH signaling upregulates genes associated with microtubule depolymerization, suggesting a transcriptional mechanism by which HH signaling influences astral microtubule dynamics and, consequently, mitotic spindle orientation. These findings highlight a novel link between a morphogen, transcriptional regulation, and the control of mitotic spindle orientation, with implications for development and tissue homeostasis.

## Introduction

The orientation of the mitotic spindle is a fundamental determinant of cell fate and tissue organization (Lancaster and Knoblich, 2012; Lu and Johnston, 2013; di Pietro et al., 2016; Bergstralh et al., 2017; Lechler and Mapelli, 2021). Proper spindle orientation ensures balanced proliferative and differentiative divisions, influencing tissue architecture and organogenesis. Spindle misorientation is implicated in developmental disorders, tumorigenesis, and tissue degeneration, underscoring the importance of precise regulatory mechanisms.

Mitotic spindle orientation is regulated by highly conserved intracellular machinery, including astral microtubules and a force-generating complex composed of the heterotrimeric Gα protein Gαi, leucine/glycine/asparagine repeat–containing protein (LGN), nuclear mitotic apparatus protein (NUMA), and dynein (Lancaster and Knoblich, 2012; Lu and Johnston, 2013; di Pietro et al., 2016; Bergstralh et al., 2017; Lechler and Mapelli, 2021). Astral microtubules grow from the centrosome toward the cell cortex, where they are captured by the Gαi–LGN–NUMA–dynein complex. Localized pulling force exerted on the astral microtubules by the force-generating complex orients the mitotic spindle (Kiyomitsu, 2019). Despite their critical roles in development and tissue homeostasis, how secreted factors regulate this intracellular machinery remains poorly understood.

Neural progenitor cells, particularly ventricular radial glia (vRGs), rely on regulated spindle orientation to balance self-renewal and differentiation (Sanada and Tsai, 2005; Postiglione et al., 2011; Das and Storey, 2012; Xie et al., 2013; Falk et al., 2017; Li et al., 2017). vRGs initially undergo symmetric divisions to expand the progenitor population; these are followed by asymmetric divisions that generate neurons. During the neurogenic phase, the orientation of the mitotic spindle of a vRG relative to the ventricular surface is highly associated with the fate of its daughter cells (Shitamukai et al., 2011; LaMonica et al., 2013). vRGs dividing with a mitotic spindle parallel to the ventricular surface (horizontal division) mostly produce neurons or intermediate progenitors, whereas those dividing obliquely or vertically (collectively termed non-horizontally) produce outer radial glia (oRGs). Importantly, the disproportionate increase in oRGs in higher mammals contributes to neocortical expansion and folding (Fietz et al., 2010; Hansen et al., 2010; Reillo et al., 2011). Therefore, elucidating the mechanisms that control the vRG division angle is crucial to understanding brain development and evolution.

Hedgehog (HH) signaling regulates cell fate determination and proliferation in multiple tissues (Briscoe and Therond, 2013). We previously demonstrated that HH signaling is a conserved mechanism that promotes non-horizontal vRG division in mice, ferrets, and human cerebral organoids (Wang et al., 2016; Hou et al., 2021). However, how HH signaling modulates vRG division angle is unknown. Here, we reveal that HH signaling alters astral microtubule dynamics through a canonical pathway. Notably, this function of HH signaling is not restricted to vRGs.

## Materials and Methods

### Mice

All animal procedures were approved by the Institutional Animal Care and Use Committee of St Jude Children’s Research Hospital. We used the following mouse strains: *SmoM2*^*flox*^ (Jackson Laboratory [JAX], 005130), *GFAP::Cre* (JAX, 004600), and *Gli2*^*flox*^ (JAX, 007926). All mice were maintained in a mixed genetic background, and all were maintained on a 12-h dark/light cycle. We used both sexes of mice for experiments. A vaginal plug is first observed in female mice on embryonic day 0.5 (E0.5).

### iPSC and human cerebral organoid culture

Induced pluripotent stem cells (iPSCs) expressing GFP-α-tubulin from the endogenous *TUBA1B* gene were obtained from the Allen Institute/Coriell (AICS-0012) and were maintained in mTeSR™1 medium (STEMCELL technologies). To investigate the effect of HH signaling on the iPSC division angle, iPSCs were seeded on µ-Slides (ibidi, 80426) and treated with 400 nM SAG (Smoothened agonist) or DMSO for 24 h before time-lapse imaging or fixation.

Cerebral organoids were generated using the protocol described previously (Lancaster et al., 2013; Wang et al., 2016; Wang et al., 2022). Briefly, to generate embryoid bodies (EBs), iPSCs were suspended in medium consisting of DMEM/F12 (Life Technologies, 11330-032) supplemented with 10% knockout serum replacement (KOSR) (Life Technologies, 10828-028), 3% ES-quality FBS (Life Technologies, 10439-016), 1% GlutaMAX (Life Technologies, 35050-061), 1% MEM-NEAA (Life Technologies, 11140-050), 7 ppm (v/v) β-mercaptoethanol (Life Technologies, 21985-023), 4 ng/mL bFGF (Peprotech, 100-18B), and 50 µM Rho-associated kinase (ROCK) inhibitor (ATCC, ACS-3030) and seeded at 9000 cells per 150 µL in each well of a 96-well Lipidure®-Coat plate (Gel Company, LCV96). The medium was changed every other day for 6 days, omitting the bFGF and ROCK inhibitor after day 4. Then, EBs were transferred to wells of a Costar® 24-well plate (Corning, 3473) (1 EB per well) and fed every other day with neural induction medium consisting of DMEM/F12 supplemented with 1% N-2 supplement (Life Technologies, 17502-048), 1% GlutaMAX, 1% MEM-NEAA, and 1 µg/mL heparin for 4–5 days until neuroepithelial morphology became evident. The neuroepithelial aggregates were then embedded in a drop (15 µL) of Matrigel (Corning, 356234). The embedded aggregates (*n*□= □16) were grown in 6-mm dishes containing 5 mL of differentiation medium (50% DMEM/F12, 50% Neurobasal Medium, 0.5% N-2 supplement, 1% B-27 supplement without vitamin A (Life Technologies, 12587-010), 0.025% (v/v) human insulin (Sigma, I9278), 3.5 ppm (v/v) β-mercaptoethanol, 1% GlutaMAX, 0.5% MEM-NEAA, and 1% antibiotics/antimycotics) with constant shaking at 75 rpm for 4 days; the medium was changed on the second day. Four days after differentiation, the tissue droplets were fed with differentiation medium containing B-27 supplement with vitamin A (Life Technologies, 17504-044) and incubated at 37°C in 5% CO_2_ with constant rotation at 75 rpm, with the medium being replenished every 3 days.

To investigate the effects of HH signaling on the vRG division angle, 5-week-old cerebral organoids were cut into 300-µm slices with a vibratome (Leica VT1200S). Slices were placed in 35-mm glass-bottom dishes (MatTek, P35G-0.170-14-C) and treated with SAG (400 nM) or DMSO for 24 h before time-lapse imaging.

### Immunostaining and microscopy

Embryonic brains were fixed overnight in 4% paraformaldehyde (PFA) in PBS, cryoprotected for 24 h in 30% sucrose in PBS at 4°C, and embedded in OCT medium (Sakura Finetek). Tissue blocks were cryosectioned at a thickness of 12 µm. iPSCs were fixed in 4% PFA for 45 min at 4°C and washed with PBS. Immunostaining was performed using primary antibodies against EB1 (Abcam, ab53358; diluted 1:1000), GFP (Proteintech, gba488; 1:200), phospho-histone H3 Ser10 (Abcam, 14955; 1:1,000), and phospho-vimentin Ser55 (MBL International, D076-3; 1:2,000). Alexa Fluor®–conjugated antibodies (Invitrogen) were used as secondary antibodies. DNA was stained with DAPI (Cayman Chemical, 14295).

Images of immunostained tissue sections were acquired with a Zeiss 780 microscope. Time-lapse images of live organoid slices and iPSCs were acquired on a 3i Marianas spinning disk confocal system with environmental control. Images were obtained at 5-min intervals.

### RNA sequencing and analysis

Total RNA was extracted using an RNeasy Mini Kit (Qiagen). Libraries were prepared using a TruSeq Stranded Total RNA Kit (Illumina) and sequenced with an Illumina HiSeq system. The raw reads were trimmed with Trim-Galore version 0.60 and mapped to the GRCh38 human genome assembly with STAR v2.7. The gene levels were then quantified with RSEM v1.31, based on GENCODE Human Release 3. Genes with low counts (CPM□<□0.1) were removed from the analysis, and only protein-encoding genes were used for differential expression analysis. Normalization factors were generated using the TMM (Trimmed Mean of M-values) method. Counts were then transformed with voom and analyzed with the lmFit and eBayes functions of R limma package version 3.42.2. The false discovery rate (FDR) was estimated using the Benjamini– Hochberg method. Pre-ranked Gene Set Enrichment Analysis (GSEA) (Subramanian et al., 2005) was performed by using the −log10(*P* value)*log2 fold change value (from differential expression analysis) ranked gene list against gene sets in the Molecular Signatures Database (MSigDB v2023.1), including the GO Molecular Function, GO Biological Process, GO Cellular Component, Reactome, CanonicalPathway, and Hallmark gene sets. GSEA version 4.3.2 was used with the following parameters: number of permutations = 1000, permutation type = gene_set, metric for ranking genes = Signal2Noise, enrichment statistic = weighted.

### Statistics

Statistical analysis was performed using GraphPad Prism software. Normality was assessed by Kolmogorov–Smirnov testing. The two-tailed, unpaired *t*-test and the Mann–Whitney test were used for normally and non-normally distributed data, respectively. The Spearman correlation test was used for the EB1 and division angle correlation analysis. The chi-square test was used to analyze the division modes of vRGs in the embryonic cortex. In each case, data were obtained from at least three biological replicates.

## Results

### HH signaling increases the vRG division angle in cerebral organoid slice culture

To investigate cellular mechanisms by which HH signaling affects the vRG division angle, we used time-lapse microscopy to observe vRGs undergoing mitosis in cerebral organoids. To visualize the mitotic spindle in live cells, we generated cerebral organoids by using iPSCs expressing green fluorescent protein (GFP)-tagged α-tubulin from the endogenous *TUBA1B* locus. We treated sections of 5-week-old organoids with SAG (an agonist of Smoothened [SMO], a key HH pathway activator) or DMSO (as a control) for 24 h and acquired time-lapse images of mitotic vRGs at 5-min intervals (Fig. 1A). Activation of HH signaling by SAG significantly increased vRG division angles (Fig. 1B). Notably, SAG increased the mitotic spindle rotation (Fig. 1C) and prolonged mitosis (Fig. 1D). These results suggest that HH signaling deters anchoring of the mitotic spindle, leading to increased spindle rotation and delayed execution of mitosis. Inhibiting HH signaling with a SMO antagonist, SANT1, did not affect the division angle (data not shown), which is consistent with observations in fetal ferret cortex slice cultures (Hou et al., 2021). In slice cultures, secreted endogenous HH ligands might be too diluted by the culture medium to affect the division angle; if so, further inhibition by SANT1 would have little or no effect.

**Figure 1.**
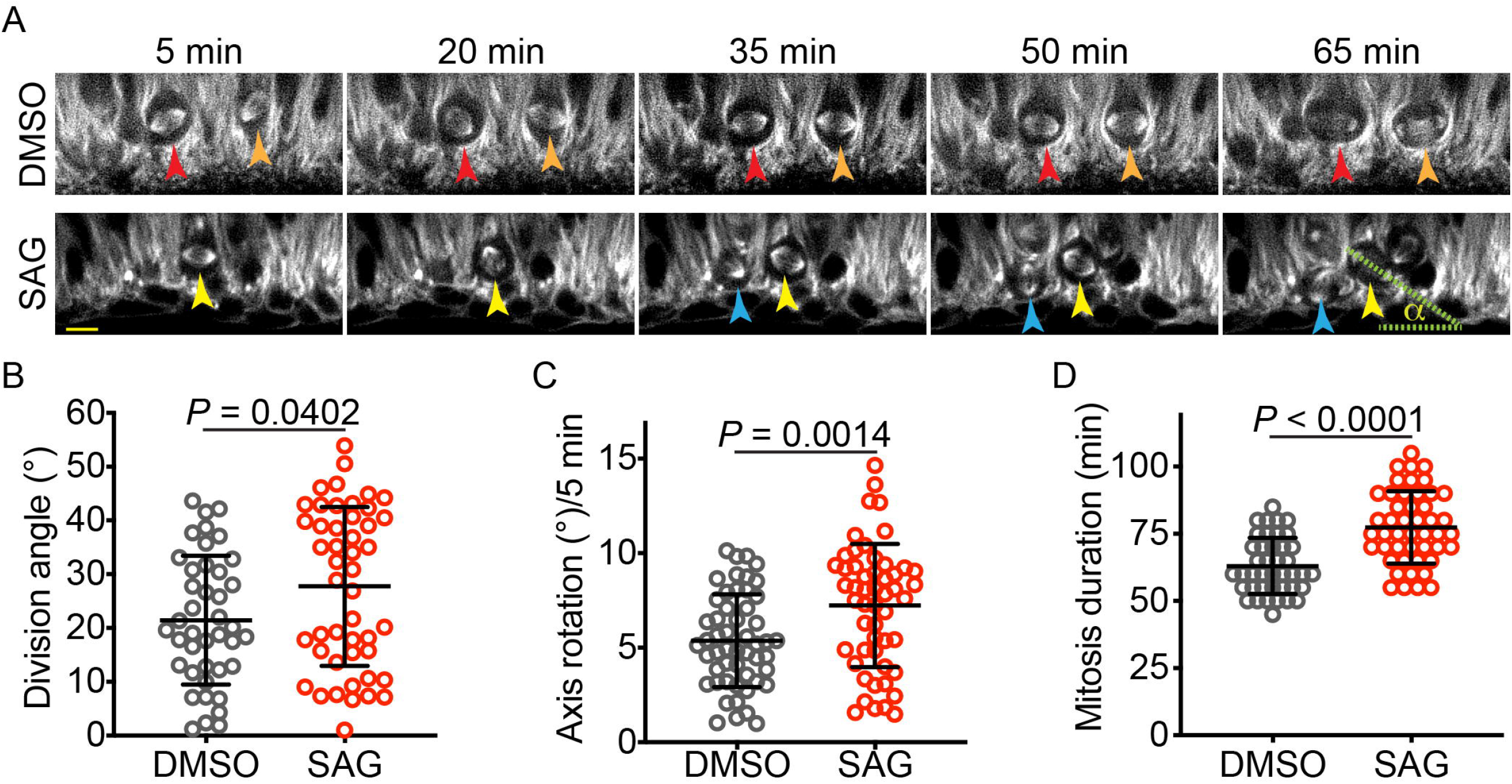
SAG increases the vRG division angle in cerebral organoid slice culture. (A) Time-lapse images of dividing vRGs expressing GFP-α-tubulin from the endogenous *TUBA1B* gene. Arrowheads of the same color track individual cells over time. “α” indicates the division angle relative to the apical surface. Scale bar: 20 μm. (B–D) Quantifications of the vRG division angle, mitotic spindle rotation rate, and mitosis duration.

### HH signaling increased the iPSC division angle in monolayer culture

To determine whether HH signaling affected mitotic spindle orientation, specifically in vRGs or more broadly in other cell types, we treated iPSCs with SAG or DMSO for 24 h before fixation and measured their division angles relative to the culture substratum. Surprisingly, SAG significantly increased the iPSC division angle (Fig. 2A, B). This observation was confirmed by live-cell imaging (Fig. 2C, D). Moreover, similar to the effects observed in vRGs, SAG increased the mitotic spindle rotation and prolonged mitosis in iPSCs (Fig. 2E–G). These results suggest that HH signaling influences mitotic spindle orientation in multiple cell types, including iPSCs.

**Figure 2.**
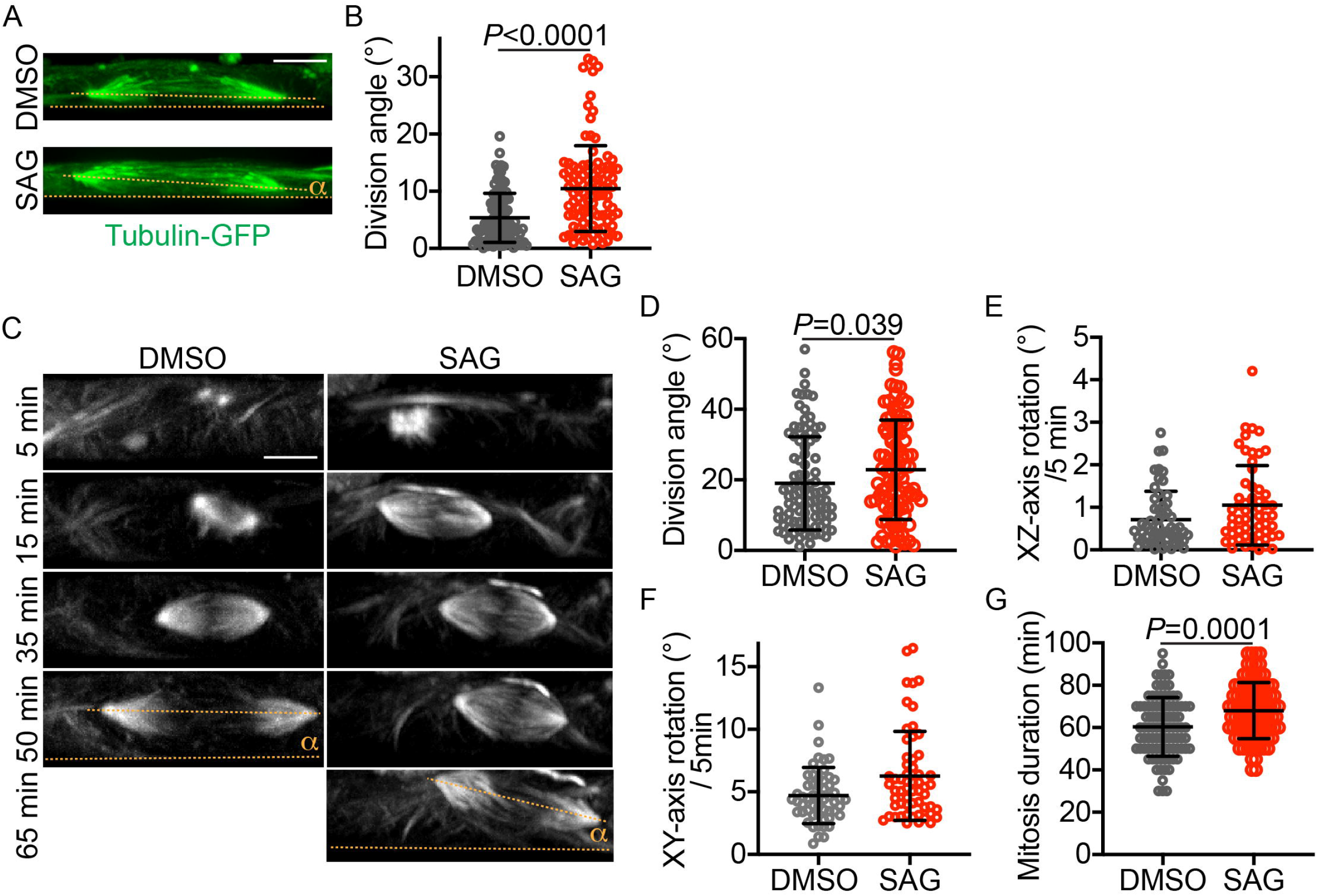
SAG increases the iPSC division angle in monolayer culture. (A) Representative XZ projections of GFP-α-tubulin–expressing iPSCs immunostained with an anti-GFP antibody. The division angle (α) is measured relative to the culture plate surface (i.e., the XZ-axis angle relative to the X-axis, indicated by dotted lines). (B) Quantification of the division angles of fixed iPSCs, as shown in panel A. (C) Time-lapse XZ projections of dividing iPSCs that express GFP-α-tubulin. (D–G) Quantifications of the iPSC division angle, mitotic spindle rotation rate, and mitosis duration. Scale bar: 5 μm.

### HH signaling reduces astral microtubule number and length

Because astral microtubules anchor the mitotic spindle to the plasma membrane and contribute to spindle positioning and orientation, we investigated the effects of HH signaling on astral microtubules. We treated iPSCs expressing GFP-α-tubulin with SAG or DMSO and visualized the astral microtubules with an anti-GFP antibody. To enable 3D tracing and quantification of astral microtubules, we developed a napari tool (napari 3D filament annotator; https://github.com/amedyukhina/napari-filament-annotator). By using this tool, we found that SAG significantly decreased both the number and the length of astral microtubules, without affecting their bending angle or curvature (Fig. 3A–E). These findings were corroborated by quantifying EB1 (end-binding protein 1, also known as microtubule-associated protein RP/EB family member 1), a protein that binds to the plus-end of microtubules (Morrison and Askham, 2001). SAG treatment significantly reduced the number of EB1-positive puncta (Fig. 3A, F). The number of EB1 puncta per cell was larger than the number of astral microtubules per cell, reflecting the stronger fluorescent intensity of the former. Interestingly, analysis of combined data from DMSO-treated and SAG-treated cells revealed a negative correlation between the number of EB1 puncta and the iPSC division angle relative to the substrate (Fig. 3G), suggesting that the reduction in astral microtubules contributes to the tilting of the mitotic spindle away from its parallel orientation relative to the substrate.

**Figure 3.**
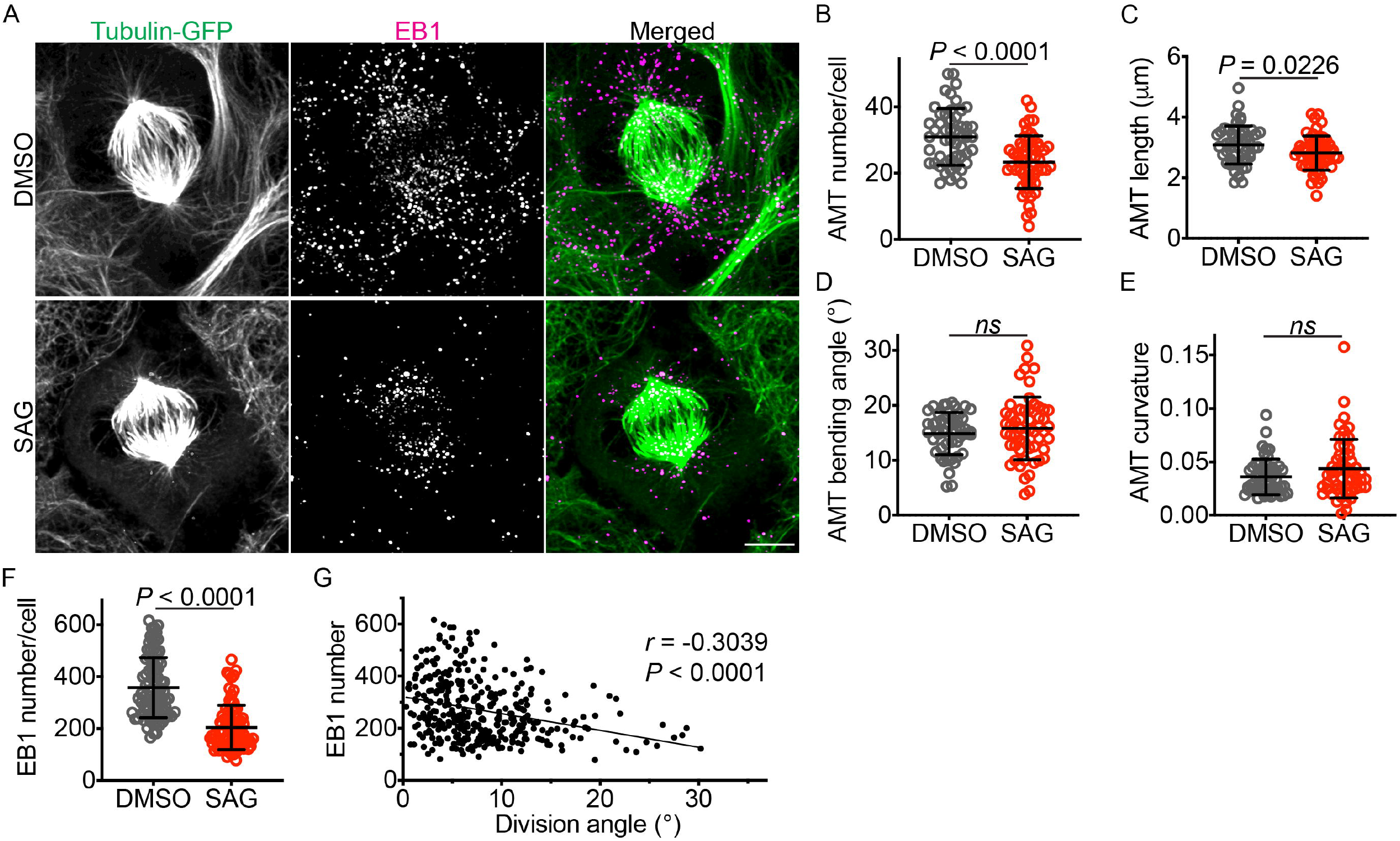
SAG decreases the number and length of astral microtubules. (A) GFP-α-tubulin–expressing iPSCs labeled with an anti-GFP antibody and an anti-EB1 antibody. Scale bar: 5 μm. (B–E) Quantifications of astral microtubule number, length, bending angle, and curvature. (F) Quantification of EB1 puncta. (G) A plot showing a negative correlation between EB1 puncta number and division angle.

We also examined whether HH signaling affected the levels and localization of NUMA and LGN, but we obtained no conclusive results.

### Canonical HH signaling modulates mitotic spindle orientation

Canonical HH signaling regulates gene expression through GLI transcription factors, whereas non-canonical HH signaling operates independently of GLI factors (Akhshi et al., 2022). Notably, non-canonical HH signaling modulates the actin cytoskeleton, which interacts with astral microtubules and regulates mitotic spindle orientation (Kunda and Baum, 2009; Kwon et al., 2015; Yu et al., 2019). To investigate whether canonical HH signaling regulated mitotic spindle orientation, we analyzed the vRG division angle in wild-type, *GFAP::Cre; SmoM2*^*fl/+*^, and *GFAP::Cre; SmoM2*^*fl/+*^,*Gli2*^*fl/fl*^ embryos at embryonic day 14.5. SMOM2 is a constitutively active form of SMO. As reported earlier (Wang et al., 2016; Hou et al., 2021), activation of HH signaling by SMOM2 expression in the *GFAP::Cre; SmoM2*^*fl/+*^ embryonic cortex significantly decreased the proportion of horizontal vRG divisions compared to that in wild-type embryos (Fig. 4A, B). Importantly, loss of GLI2 in *GFAP::Cre; SmoM2*^*fl/+*^,*Gli2*^*fl/fl*^ embryos partially restored the proportion of horizontal vRG divisions (Fig. 4B), demonstrating that HH signaling modulates the vRG division angle, at least in part, through the canonical, GLI-dependent pathway.

**Figure 4.**
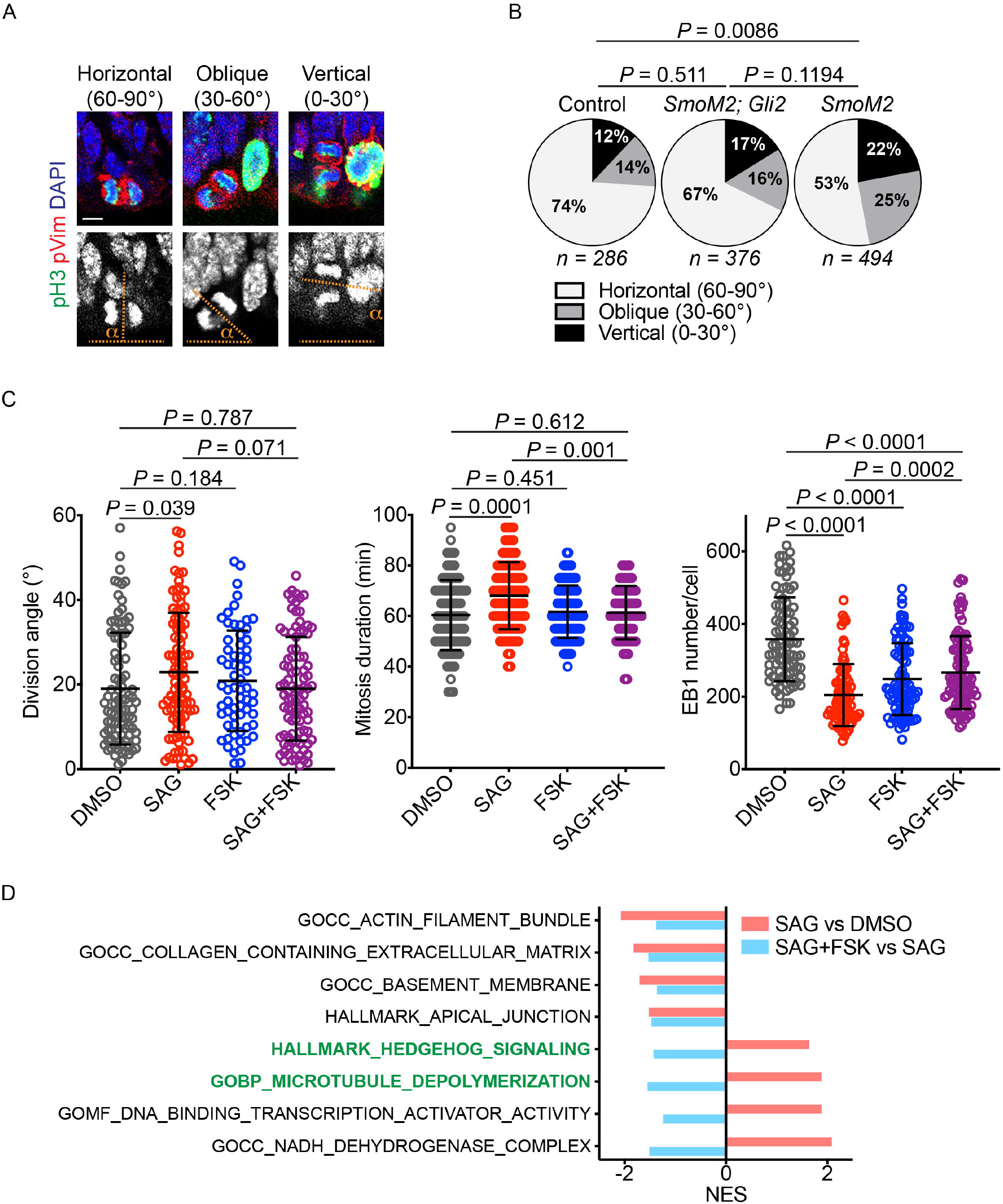
HH signaling modulates the division angle through the canonical pathway. (A) Mitotic vRGs in E14.5 mouse embryonic cortex immunostained for phospho-histone 3 (pH3, green), phospho-vimentin (pVim, red), and DAPI (blue and gray in the upper and lower panels, respectively). pH3 marks mitotic cells, and pVim visualizes the division plane. Scale bar: 5 μm. (B) Quantification of the iPSC division angle, mitotic duration, and EB1 puncta number. Data for the DMSO and SAG groups are reproduced from Figures 2 and 3. (D) Normalized enrichment scores (NESs) for gene sets enriched in iPSCs treated with SAG vs. DMSO and with SAG+FSK vs. SAG.

We then investigated whether canonical HH signaling regulated the iPSC division angle. Because protein kinase A (PKA) is a conserved key inhibitor of active GLI transcription factor formation (Wang et al., 1999; Wang et al., 2000), we treated iPSCs with forskolin (FSK), a PKA activator, to block SAG-mediated activation of canonical HH signaling. Consistent with the results obtained in mouse embryos, FSK reversed the effects of SAG on the division angle, the duration of mitosis, and the number of EB1 puncta (Fig. 4C). These results indicate that HH signaling modulates astral microtubules and mitotic spindle orientation, at least in part, via the canonical pathway that regulates gene expression.

To further elucidate how HH signaling regulated mitotic spindle orientation, we performed RNA sequencing and GSEA on iPSCs treated with DMSO, SAG, or SAG+FSK (Fig. 4D, Supplementary Figure S1). As expected, SAG increased the expression of the HH signaling gene set, and this effect was reversed by FSK. SAG also upregulated a gene set associated with microtubule depolymerization, and this upregulation was also reversed by FSK. Additionally, SAG decreased the expression of gene sets involved in the actin filament, extracellular matrix, basement membrane, and apical junction, all of which can influence mitotic spindle orientation and positioning (Thery et al., 2005; Toyoshima and Nishida, 2007; Kunda and Baum, 2009; Fink et al., 2011; Kwon et al., 2015; Yu et al., 2019; Lechler and Mapelli, 2021; Naher et al., 2025). However, negative enrichments of these gene sets were not reversed by FSK, unlike those of the HH signaling and microtubule depolymerization gene sets. These data suggest that HH signaling decreases the number and length of astral microtubules, at least in part, by upregulating genes involved in microtubule depolymerization.

## Discussion

Our findings have revealed a novel role for HH signaling in modulating mitotic division dynamics. We have demonstrated that increased HH signaling leads to a reduction in both the length and the number of astral microtubules, increased mitotic spindle rotation, prolonged mitosis, and a higher incidence of non-horizontal divisions. The observed decrease in astral microtubule length and number provides a mechanistic link to the other phenotypic changes. The reduction in astral microtubules, which are crucial for anchoring the spindle to the cell cortex, could directly contribute to the increased spindle rotation. This increased rotation may, in turn, lead to non-horizontal anchoring of the spindle pole. Furthermore, the reduced astral microtubule network may delay the stable anchoring of the spindle pole, thus prolonging mitosis. Our data are consistent with previous work in mouse vRGs, which showed that non-horizontally dividing cells exhibit increased spindle rotation and extended mitosis (Haydar et al., 2003) and that these cells also exhibit fewer astral microtubules when compared with horizontally dividing vRGs (Mora-Bermudez et al., 2014). These results suggest that HH signaling influences the geometry and timing of vRG division, potentially affecting the overall production and organization of cortical neurons. Notably, the similar influence of HH signaling on mitotic spindle orientation in both vRGs and iPSCs suggests that spindle orientation serves as a conserved mechanism through which HH signaling regulates development and homeostasis.

The control of mitotic spindle orientation is a fundamental mechanism across species and cell types for determining daughter cell fate. Although extensive research has established the importance in this process of conserved intracellular machinery—such as astral microtubules and the Gαi–LGN–NUMA–dynein complex—and has revealed the regulatory roles of post-translational modifications and protein–protein interactions (Lancaster and Knoblich, 2012; Lu and Johnston, 2013; di Pietro et al., 2016; Bergstralh et al., 2017; Lechler and Mapelli, 2021), the influence of secreted factors remains relatively understudied, despite their key roles in development and tissue homeostasis. Our findings highlight the need for studies exploring the role of secreted extracellular cues, such as morphogens, and transcriptional regulation in mitotic spindle orientation. Further investigation is required to determine the precise molecular mechanisms by which HH signaling regulates astral microtubule dynamics.

## Acknowledgments

We thank the staff of the Animal Resource Center, the Hartwell Center for Bioinformatics and Biotechnology, and the Cell and Tissue Imaging Center at St. Jude Children’s Research Hospital for technical assistance. We thank Keith A. Laycock, PhD, ELS, for scientific editing of the manuscript. This work was supported by the American Lebanese Syrian Associated Charities (ALSAC) and by the National Institutes of Health (R01NS100939 to Y.-G. H.). The content is solely the responsibility of the authors and does not necessarily represent the official views of the National Institutes of Health.

