## Supplementary figures and images for "Canonical Hedgehog Signaling Controls Astral Microtubules and Mitotic Spindle Orientation in Neural Progenitors and iPSCs"

### Supplemental Figure 1

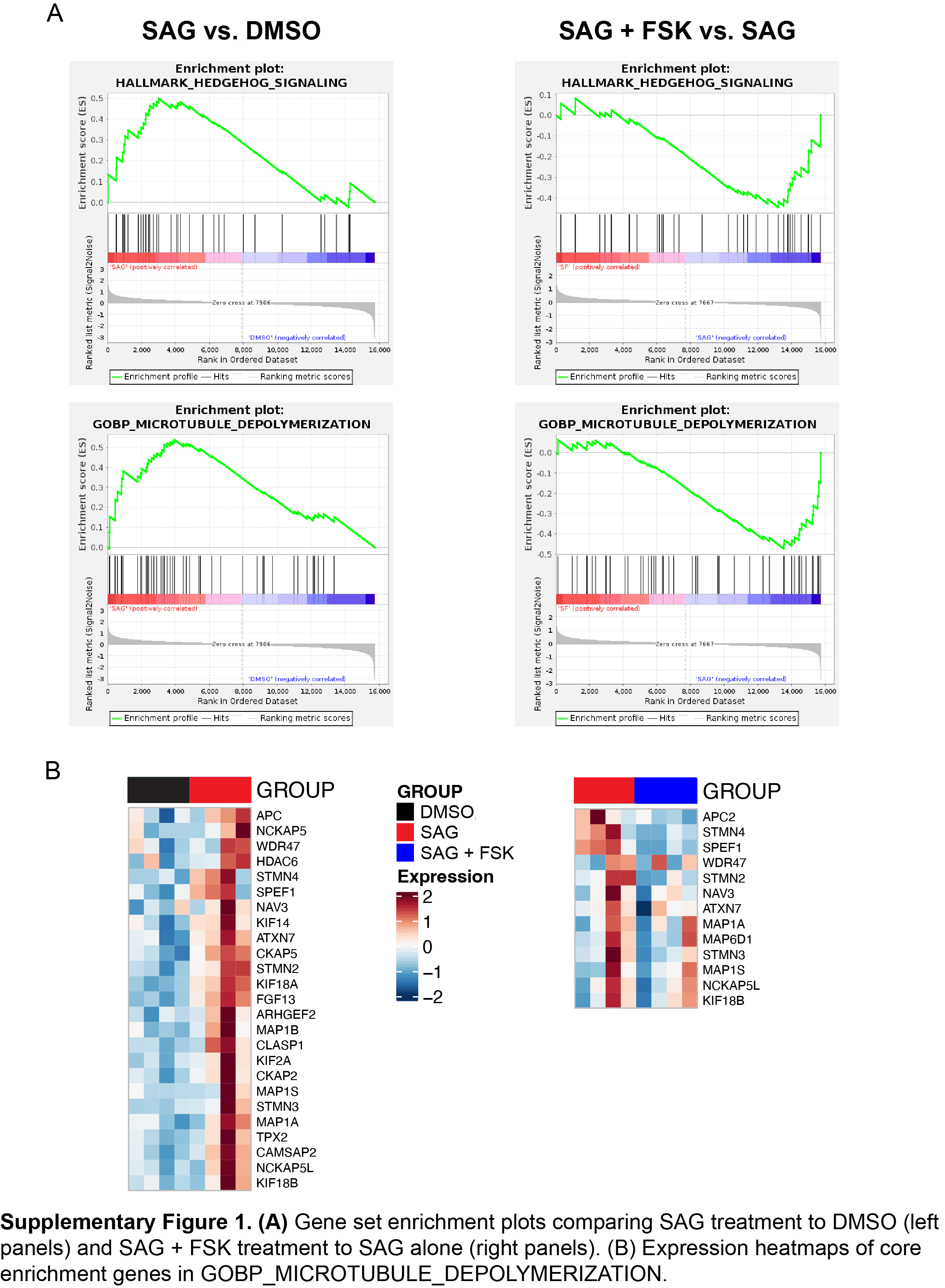
